# Improving chromatin-interaction prediction using single-cell open-chromatin profile and making insight about the cis-regulatory landscape of the human brain

**DOI:** 10.1101/2020.12.29.424732

**Authors:** Neetesh Pandey, Omkar Chandra, Shreya Mishra, Vibhor Kumar

## Abstract

Single-cell open-chromatin profiles have the potential to reveal the pattern of chromatin-interaction in a cell-type. However, currently available cis-regulatory network prediction methods using single-cell open-chromatin profiles focus more on local chromatin-interactions despite the fact that long-range interaction among genomic sites plays a significant role in gene regulation. Here, we propose a method that predicts both local and long-range interactions among genomic sites using single-cell open chromatin profiles. Using our method’s better sensitivity, we could predict almost 0.7 million interactions among genomic sites across 7 cell-types in the human brain. The chromatin-interactions estimated in the human brain revealed surprising but useful insight about target genes of human-accelerated-elements and disease-associated mutations.

## Introduction

Spatial interactions between different genome loci are required for multiple regulatory functions(1). Many groups have profiled chromatin-interaction in multiple cell types using different experimental high-throughput methods to study such complex patterns in chromatin architecture and gene regulation. The experimental methods based on chromosome conformation capture (3C) are more focused on local genomic loci (2). The ChiA-PET (Chromatin-interaction Analysis by Paired-End Tag Sequencing) method captures distal interactions, but it is limited to only binding sites of the protein of interest (3). The HIC (high-throughput chromosome conformation capture) assay provides a genome-wide chromatin-interaction profile but requires deep-sequencing to achieve high resolution(1).

Several groups have recently attempted to predict chromatin-interactions using linear one dimensional genetic and epigenetic information (4). Most of the tools proposed for predicting interaction use epigenetic information from bulk samples, often consisting of multiple cell-types (5). Simultaneous availability of many epigenome profiles is currently possible for only few cell-types. Hence predicting cell-type-specific chromatin-interaction is not trivial for many cell-types. On the other hand, if we exploit heterogeneity in the activity of genomic sites in single-cells, we could predict chromatin-interactions in a cell-type. Especially for understanding regulatory mechanism in minor cell-types in heterogeneous clinical samples for personalized therapy, single-cell epigenome profiles can provide the landscape of activity of genomic sites as well as prediction of chromatin-interaction. With experimental assays like 3C and HiC and computational methods using bulk epigenome profiles, it would not be trivial to profile chromatin-interaction maps for multiple cell-types for heterogeneous clinical samples from patients on a regular basis. Recently, Pliner et al. (6) proposed a method called Cicero to predict local chromatin-interaction using scATAC-seq (single-cell Assay for Transposase-Accessible Chromatin using sequencing) profile. However, Cicero is not sensitive in predicting distal interactions among genomic sites which are more than 500 Kbp (kilobase pairs) apart. Another method called as JRIM (jointly reconstruct *cis*-regulatory interaction maps) (7) uses open chromatin profiles of multiple cell-types to infer reliable chromatin-interactions, hence it is of less use for prediction for a single cell-type. JRIM is also designed to predict chromatin-interactions with in 500 kb window. However, it has been shown before that mutations identified by genome wide association studies (GWAS) could be influencing genes lying more than 500 Kb away. The median size of topologically associated domain (TAD) in mouse cells have been reported to 880kbp(8). Hence, predicting distal and inter chromosomal interactions using single cell epigenome profile is still an open problem of high utility. Here we developed a method called as single-cell Epigenome based chromatin-interaction Analysis (scEChIA) which can predict interactions among distal sites with high accuracy using single-cell open-chromatin profiles. We have further shown its utility in prediction of chromatin-interactions in brain cells for making useful insights.

## Result

Our computational approach is based on well-known pattern of DNA looping and chromatin conformation. Previously Tang et al. have shown that in spite of many cell-type specific interactions, multiple chromatin-interactions show high similarity across different cell-type. It is known that CTCF mediated chromatin-interaction and looping are mostly conserved and have major impact on chromatin architecture. Similarly, many short tandem repeat define boundaries of topologically associated domain (TADs) which tend to be conserved across different cell types (9). Hence, unlike Cicero, we used pre-existing knowledge of chromatin-interactions in multiple cell-types as a prior while estimating the Gaussian graphical model, using L1 regularization to predict chromatin-interaction using single cell open-chromatin profile. For this purpose, we use average value of enrichment of known chromatin-interactions in multiple cell-types to calculate L1 regularization (ρ) parameter. In addition to using sensitive L1 normalization, scEChIA uses its inbuilt function for matrix factorization to reduce noise in read-count matrix to further improve the accuracy of prediction of chromatin-interaction (see supplementary methods).

### scEChIA improves sensitivity for predicting distal interactions with high accuracy

We compared our method’s accuracy and sensitivity with the famous method Cicero (6). For this purpose, we used scATAC-seq profiles of K562, GM12878 and H1ESC published by Buenrostro et al. (10). We calculated the regularisation parameter ρ in graphical Lasso (Glasso) model using average of known chromatin-interaction in other cell-types for predicting chromatin-interaction for a cell-type. For example, for predicting chromatin-interaction in K562 cells we used prior (or regularisation parameter ρ) estimated using the average of HiC profile of GM12878 and hESC cells(11). For GM12878 we used scATAC-seq profile published by Buenrostro et al. and average of HiC profile of K562 and hESC to calculate regularisation parameter. We performed evaluation using HiC based enriched chromatin-interaction profile of relevant cell-types (see supplementary methods). We found that scEChIA is as accurate as Cicero, whereas using constant ρ value with Glasso did not provide comparable accuracy in predicting chromatin-interaction (Figure 1a). However, in comparison to Cicero, scEChIA predicts a substantially higher number (almost 100 times) of distal interactions with a gap of more than 500kb among interacting sites (Figure 1b). Further analysis revealed that genes with highest relative number of chromatin-interaction at their promoters also had higher expression in respective cell-type whose scATAC-seq profile was used for prediction (Figure 1C). Thus, it also proves that scEChIA can predict cell-type-specific interactions of gene promoters with other genomic locations. (see Figure 1c, supplementary Figure-1, supplementary method).

**Figure 1:**
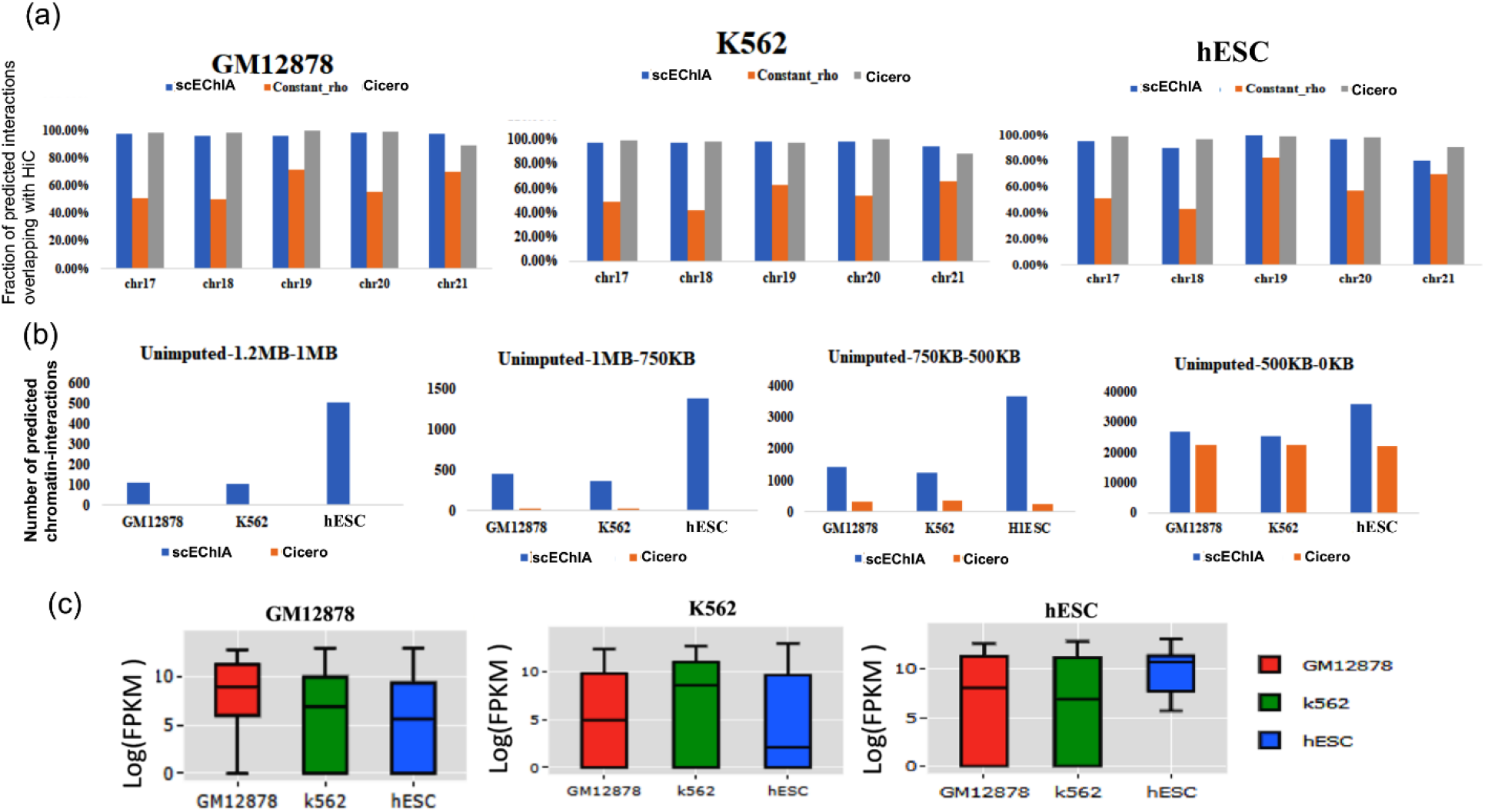
Evaluation of accuracy and sensitivity of chromatin-interaction prediction. (a) Accuracy of prediction of chromatin-interaction shown for Glasso used with constant regularisation parameter (constant_rho), Cicero and scEChIA. Accuracy was measured as the fraction of predicted chromatin-interactions, which overlapped with enriched interactions in HiC profile of respective cell (see supplementary methods). (b) The sensitivity of prediction of distal chromatin-interaction is shown here for different ranges of distance between interacting genomic loci. The sensitivity of Cicero for long-range chromatin-interactions (> 500kbp) is almost 100 times lower than scEChIA (c) Evaluation of prediction of cell-type-specific cisregulation in different by scEChIA. The expression values in terms of FPKM (fragment per kilobase per million) is shown for top 50 genes with highest relative interactions in the respective cell-type of the figure panel. Such as for panel with label GM12878, top 50 genes with highest relative number of chormation-interaction in GM12878 cells (compared to K562 and hESC) were chosen and their expression values in 3 cell types (GM12878, K562, hESC) are shown as box-plot. scEChIA can predict cell-type-specific cis-regulatory interactions. Therefore, genes with such cell-type-specific interactions also have higher expression in target cells relative to other cell-types.

### The chromatin-interaction landscape of the human brain

Recently, Lake et al. (12) published single-cell RNA-seq and scATAC-seq profile of cells derived from the adult human brain. We used scEChIA to predict chromatin-interaction in 7 brain cell-types using scATAC-seq profile published by Lake et al(12). The cell types for which we predicted chromatin-interaction are inhibitory neurons, excitatory neurons, Astrocytes, Oligodendrocytes, Oligodendrocyte precursor, Microglia, Endothelial cells. The number of predicted chromatin-interaction in different cell types ranged from 188857 in Microglia to 25838 in oligodendrocytes (total ~0.7 million interactions) (see supplementary table −1). We used HiC profile for astrocytes, published by ENCODE consortium. The accuracy of scEChIA for astrocytes cells was also high as 85-95% of predicted interactions overlapped with top interacting-loci in HiC profile in most of the chromosomes (see supplementary Figure-2). Such accuracy of scEChIA for astrocytes also hints about the reliability of predicted interactions in other six brain cell-types. Intersecting our predicted chromatin-interaction with available expression quantitative trait locus (eQTL) in the brain (13) using PGLtool (14) revealed possible cell-type in which the eQTLs are connected to their target genes. In the absence of availability of chromatin-interaction in brain cells it is not trivial to retrieve information about possible cell-type for the action of the published brain eQTLs. The results of the intersection of eQTL data-set and predicted chromatin interaction in 7 brain cell-types are provided in supplemental file-1.

#### Coverage of GWAS mutations and cell type specificity

Lake et al. investigated the enrichment of open chromatin signal within 100kbp around GWAS SNPs to estimate regions’ cell-type specificity associated with mental disorder. However, they did not find the target gene of GWAS SNP. Using our data-set of predicted chromatin-interaction in seven brain cell-types, we found target genes of GWAS mutations associated with mental disorders. We label a gene as target only when the 25kb genomic bin containing its promoter is interacting with bin containing the GWAS mutation. We further compared the enrichment of mental disorders with GWAS loci overlapping with sites interacting directly with a gene. Enrichment was calculated by normalization with the fraction of GWAS SNP of non-brain disorder overlapping with sites interacting with promoters (promoter-connected). We used GWAS mutations associated with non-brain disorders namely; Ulcerative colitis, lung cancer, breast cancer, bladder cancer, hepatitis A, hepatitis C, disease-waist to hip, disease-thorasic to hip, platelet count, bone mineral density, lung adenocarcinoma, lung disease severity in cystic fibrosis) in the null model to calculate enrichment. The higher enrichment of risk variants of few mental disorders compared to the null model showed cell-type-specificity in connectivity to promoters, which corroborated with previous reports (Figure 2a). Such as Alzheimer’s disease risk variants had higher enrichment in promoter-connected regions in microglia (Figure 2a). It has been reported that microglia signature genes have higher activity in the cortex on the development of late-onset Alzheimer disease(15). For schizophrenia, we found the highest enrichments of risk variants in promoter-connected regions in Excitatory Neurons (Figure 2a). Previously a few studies have shown an association of schizophrenia with a type of excitatory neurons known as pyramidal neurons(16). Figure 2a shows the enrichment of disease-associated variants in the promoter-connected regions in 7 types of cells in the brain.

**Figure 2:**
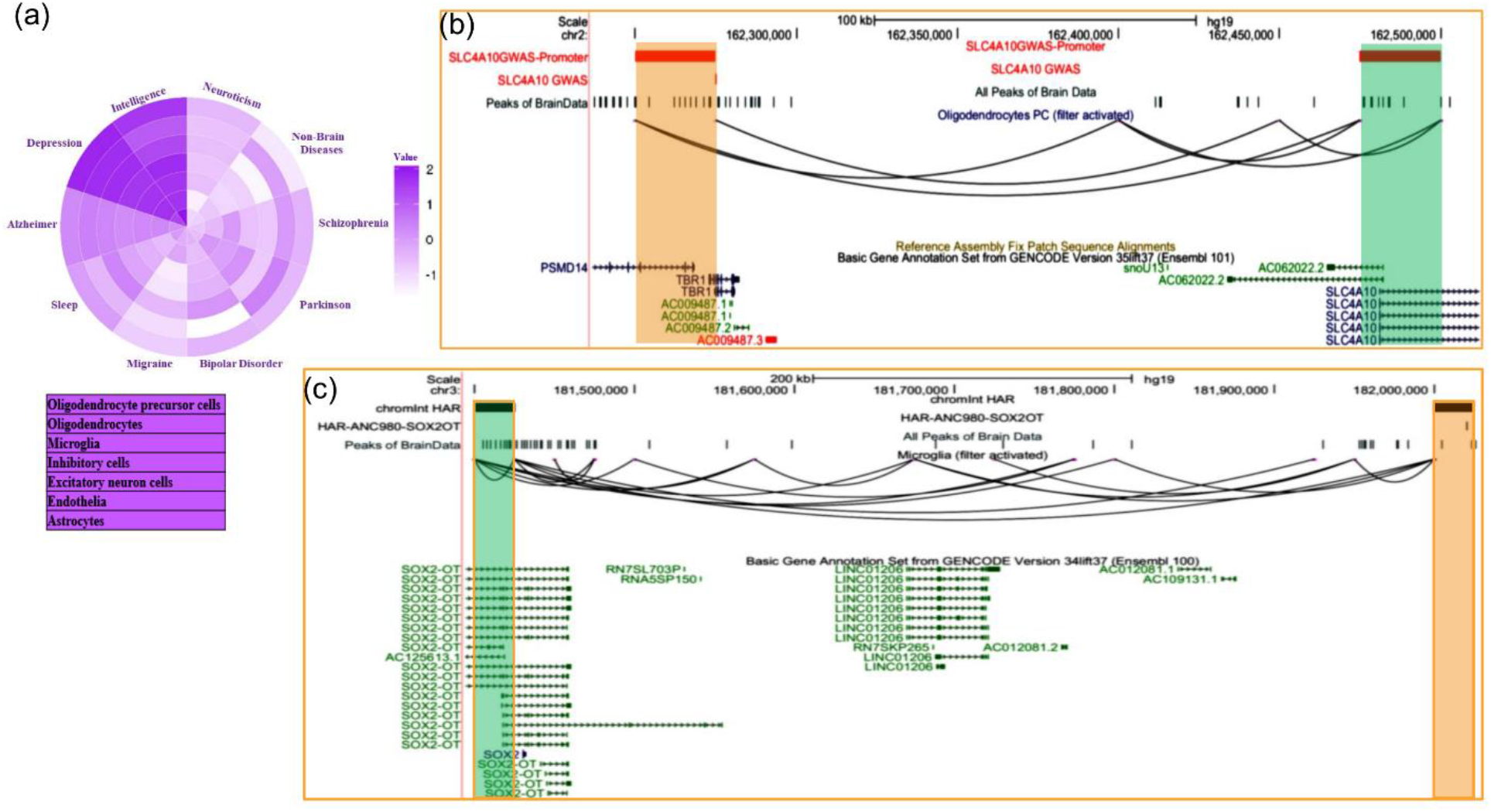
Inference about target genes for disease-associated mutations and Human accelerated elements in Brain cells. (a) Enrichment of interaction among GWAS loci and gene promoters in 7 brain cell-type for mental disorder (b) The UCSC browser snapshot showing estimated chromatin-interaction between region with GWAS SNP(rs116175783) and promoter of SLC4A10 gene. The GWAS SNP (rs116175783) is lying on intron of TBR1 gene and is associated with intelligence just as SLC4A10. (c) The UCSC browser snapshot, showing an interaction estimated to exist between a genomic bin containing a human-accelerated-region (HAR) and promoter of SOX2OT.

Our analysis also revealed a few genes with unknown association with mental disorders. While for others it revealed the possible cell-type involved in disease development through the gene. Such as a region containing a mutation (SNP id: rs116175783) associated with intelligence(17) and lying on intron of TBR1 gene appear to be interacting with promoter of gene SLC4A10 in Oligodendrocyte precursor (Figure 2b). Interestingly SLC4A10 is also known to be associated with intellectual disability (mental retardation)(18), however it’s link with SNP rs116175783 is not known, especially in oligodendrocyte precursor cells. More such results can be seen in supplementary figure 3 and supplemental file 2.

#### Target of human accelerated regions in different brain cell types

Multiple Human accelerated regions (HARS) have been discovered; however, the mechanism of effect and influence is known only for a few HARS(19). Given the fact that humans have more complex Brain structure than other species, our prediction could be a valuable resource to find target genes for HARS in brain cells. Hence, we intersected genomic sites involved in predicted chromatin-interaction with known HARS(19). Our analysis revealed several target genes for HARS, provided in the supplemental file-3 (see supplementary Figure-4). Such as scEChIA predicted interaction between a HAR named as ANC980 and promoter of gene SOX2OT (Figure 2c). SOX2OT is known to have multiple transcription-start-sites (20) and a role in the regulation of expression of SOX2 and neurogenesis. Our finding reveals that one of the promoters of SOX2OT could be regulated by a HAR, and it could have a human specific mechanism of controlling brain architecture and function.

## Discussion

Our adaptive L1 normalization approach on Gaussian graphical model and noise reduction in scATAC-seq read-count matrix predicts a higher number of distal interactions than existing methods. The predicted chromatin-interaction in 7 brain cell type could be a valuable resource for researchers to understand regulation in human brain. Especially for cells in the natural state from the in vivo brain sample, the chromatin-interaction profile availability is rare. Using our tool and predictions, one can make multiple inferences such as: cell-type specificity of the target of GWAS loci, novel associations between genes and alternative promoters with diseases, targets of HARS and alternative splicing due to cis-regulation.

## Material and Methods

### Pre-processing of Data

Our tool first divides the genome in to bins of required size. By default it uses bin of size 25kbp. For a read-count matrix it merges the peaks lying within same bin. For merging two peaks it add their read-counts. After merging the peaks, it takes log transformation of new read-count matrix as

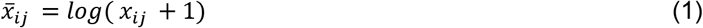

## Graphical Lasso with a penalizing parameter to embed previous knowledge

In the read-count matrix of single-cell open-chromatin profile, the number of peaks is often more than cells. Hence the estimation of matrix with covariances of peak-activity is not trivial. For such problems, Graphical Lasso(21) method helps in estimating regularized covariance matrix and its inverse. The inverse of covariance matrix can be used to calculate partial correlations. Here partial correlation provides the degree of co-accessibility between peaks, after removing the effect of confounding factor due to other peaks. Graphical Lasso is used to detect such direct association among variables. The penalty term in Graphical lasso causes shrinkage of partial correlations between peaks pairs(21), when the strength of their association is low. The Graphical Lasso method tries to minimize:

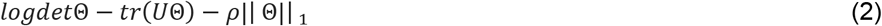

Where Θ is the inverse covariance matrix and *U* is the covariance matrix and ρ is the penalty term for L1 norm based regularization. The penalty term can be a matrix consisting of different ρ values for each pair of variables (Peaks). Our method uses a penalty matrix which is designed differently based on the knowledge of pre-existing chromatin-interaction profile. The elements of the penalty matrix are calculated as

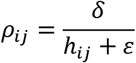

Where *h_ij_* is the average enrichment level of chromatin-interaction between genomic bins *i and j* estimated using published HiC profile of multiple cell-types. The term ε stand for a pseudo-count to stop the inflation of penalty terms in case no chromatin-interaction is found in the available HiC profiles. Whereas *δ* is a constant which can be adjusted to increase or decrease the number of predicted interactions at the cost of accuracy. The design of our method is also meant to handle the following cases

1. When two interacting sites have high activity in all cell-types, and the drop-out in their read-counts is due to random effect, then the covariance between them might be low. However, if their interaction is present in all cell types, giving a lower penalty or higher prior value would help retrieve that information.
2. If the noise level in read-counts of single-cell open chromatin profile is high then a prior guess about the background could lead to an improved prediction of interaction.
3. If two sites have cell-type-specific interactions and show decent co-accessibility, it could still be retrieved as the penalty is not exponentially high.
4. If two sites lying far away from each other have high covariance, it could be due to cis or trans effect (indirect interaction). Trans effect can be taken care of to some extent by calculating partial correlation, as well as using prior knowledge that they have low chances of direct interaction.

Hence prior knowledge (or guess of penalty matrix) is a crucial step. In order to further improve the prediction, scEChIA uses matrix factorization to reduce noise in the read-count. The matrix-factorization used by scEChIA is described in Supplementary methods.

## Program availability

The scEChIA tool is available at https://reggenlab.github.io/scEChIA/ and http://reggen.iiitd.edu.in:1207/scEChIA/

## Conflict of Interest

The authors declare no conflict of interest.

## Notes

### Competing Interest Statement

The authors have declared no competing interest.

